# Platelets Link Coagulation and Complement in Regulating Placental Vascular Development

**DOI:** 10.1101/2025.09.11.675525

**Authors:** Arno Fenske, Lisa Schumann, Ulrike Peters-Bernard, Kerstin Flächsig-Schulz, Emma Arndt, Andreas Klos, Bryan Paul Morgan, Wioleta M. Zelek, Andreas Tiede, Markus Abeln

## Abstract

During early pregnancy, maternal blood surrounds the embryo before the placenta is fully developed, requiring tight regulation of maternal blood flow into the placental vasculature. We identify placental microthrombi (PMTs) as essential structures guiding this process. PMTs contain platelets, coagulation factors, and complement proteins, and their formation depends on maternal platelet activation by thrombin through the protease-activated receptor PAR4. Deficiency of PAR4 abolished PMTs and caused excessive bleeding at the implantation site. C3 deficiency also led to increased bleeding events, indicating that complement activation contributes to thrombosis in the placental circulation. Conversely, dysregulated complement activation in CMP-sialic acid synthase–deficient (*Cmas*^−/–^) mice led to widespread thrombosis and failed placental development. Strikingly, platelet activation via PAR4 was necessary to localize complement activation to trophoblast surfaces, thereby coupling coagulation and complement in PMT formation. Depletion of maternal platelets mitigated complement-driven thromboinflammation in *Cmas*^−/–^ pregnancies, restoring placental growth. These findings uncover a critical cooperation between platelets, coagulation, and complement in establishing maternal blood flow to the placenta. Successful pregnancy therefore requires not only activation but also tight regulation of these systems to balance necessary PMT formation with the prevention of pathological thrombosis.

**Graphical abstract:** (Created in BioRender. Tiede, A. (2025) https://BioRender.com/l6zeezr)

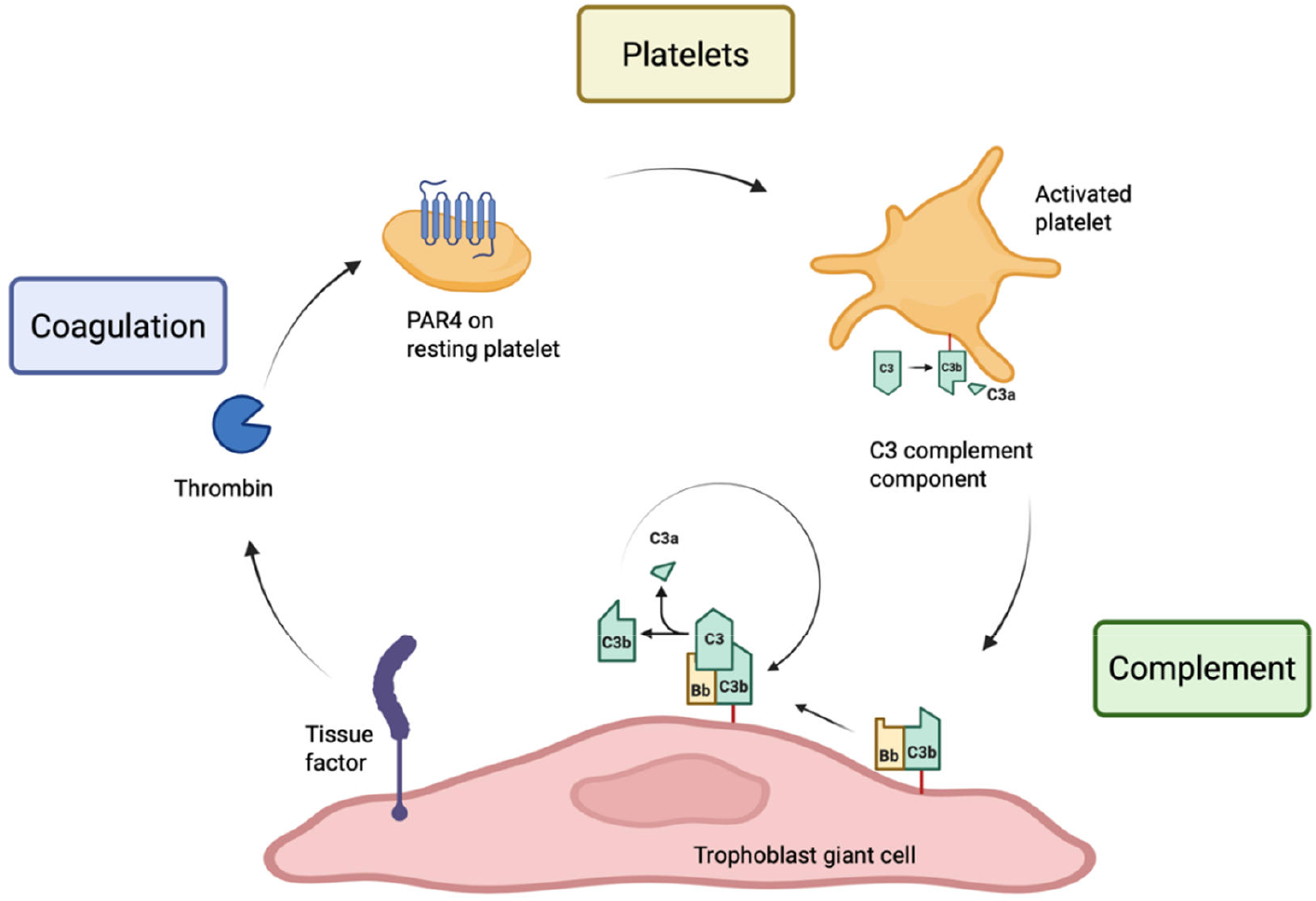

## Introduction

For a pregnancy to be successful, several challenges must be overcome, some of which appear to be mutually exclusive. On the one hand, the maternal and fetal blood must be in close proximity to ensure the foetus is adequately supplied with oxygen and nutrients; on the other hand, immunological shielding is essential to prevent an immune response against paternal antigens from the maternal immune system. Both mice and humans develop a haemochorial placenta, in which maternal blood is in direct contact with fetal trophoblasts thus enabling efficient supply of fetal tissues. This requires a variety of local and systemic immunological adaptations to allow close interaction of mother and embryo.

In the early stages of haemochorial pregnancy, before fetal blood circulation has developed, the embryo is surrounded by maternal blood. Even at this early stage, nutrients are actively transported toward the embryo by the extraembryonic tissue (1). However, rapid embryonic growth soon exceeds the capacity of this supply system. A recently published study showed that, by embryonic day 9 (E9), the implantation site becomes severely hypoxic (2). During this period, the embryonic circulation develops, and from E10 onwards, nutrient exchange between the fetal and maternal blood begins in the placenta (3). Maternal blood flow through the developing placenta must be tightly regulated and mechanisms of haemostasis and thrombosis appear to be involved in this regulation.

Trophoblast giant cells (TGCs), which form the outermost layer of the placenta at this stage, express potent procoagulant molecules like tissue factor (TF) (4), as well as anticoagulant proteins including endothelial protein C receptor (EPCR) and thrombomodulin (TM) (4). Loss of TF, EPCR, or TM are all embryonically lethal around E9, highlighting the importance of a balanced pro- and anticoagulant system. Notably, embryonic lethality caused by loss of EPCR or TM can be partially rescued by re-balancing the mechanisms of thrombosis and haemostasis, for example by inhibiting other molecules involved in thrombin generation (e.g., factor VIII [FVIII]), or maternal platelet activation via deletion of the protease activated receptor PAR4 (5, 6). Together, these findings demonstrate that while TGCs have pronounced procoagulant properties of vital importance, excessive thrombosis must be prevented at the same time.

Interestingly, deficiencies in complement regulation produce phenotypes similar to those observed in TM- and EPCR-deficient animals. Loss of the complement receptor type 1-related gene Y protein (Crry) or of CMP-sialic acid synthase (CMAS) – the latter being essential for the biosynthesis of sialic acid-bearing glycans, which enables fully functional complement regulator factor H – lead to severe deficits in placental development and ultimately embryonic death (7-9). In both models, cellular damage is mostly not driven by the formation of the membrane attack complex and the precise cause of placental malformation remains unclear (7, 9). The temporal congruence of the phenotypes of mice with deficits in anticoagulation (TM and EPCR) and complement regulation (Crry and CMAS) suggests a close link between complement, coagulation, and the developing placenta.

In this study, we examined mouse models with defects in the coagulation or complement cascades, as well as platelet activation, to explore mechanisms of thrombosis and haemostasis that are essential for successful placenta development. Our data show that PAR4-mediated activation of platelets is essential for placental microthrombus (PMT) formation at the fetal-maternal interface. The absence of PMTs leads to severe bleeding events at mid-gestation. Proteins of the complement system are also present in these placental thrombi and absence of C3 leads to an increased placental bleeding. Using a complement-sensitive mouse model (CMAS-negative), we demonstrate that excessive complement activation results in increased thrombosis, a process very similar to the pathogenic process of microvascular *thromboinflammation* described in a diverse range of human diseases (10). Notably, loss of PAR4 diminished complement deposition on CMAS-negative trophoblasts but did not rescue development of the placenta. Only severe reduction of the platelet count during pregnancy prevented complement-mediated thrombosis in CMAS-deficient pregnancies, suggesting that PAR4-negative platelets were activated by other mechanisms at the fetal-maternal interface. Together our findings demonstrate that platelet and complement activation are crucial for preventing haemorrhage during pregnancy; however, platelet-mediated complement activation needs to be tightly controlled to prevent thrombosis and severe placental pathology.

## Results

### Microthrombosis and bleeding in the developing placenta

In order for maternal blood to stop flowing around the implantation site and instead be directed through the blood vessels forming in the developing placenta, the blood flow must be blocked locally. Analysis of the boundary layer between trophoblasts and decidua on embryonic day 8.5 (E8.5) revealed the presence of placental micro thrombi (PMTs) in the vicinity of trophoblast giant cells (TGCs) on the mesometrial side of the placenta (Fig. 1A). These PMTs contained coagulation proteins (e.g., FVIII), maternal platelets (detected through platelet factor PF4 and β3 Integrin (β3)) and urokinase-type plasminogen activator receptor (uPAR), which is part of the fibrinolytic system. Interestingly, PMTs contain not only platelets and coagulation proteins, but also the complement protein C3. This is shown by reactivity for the C3 activation product C3d (Fig. 1A).

**Figure 1:**
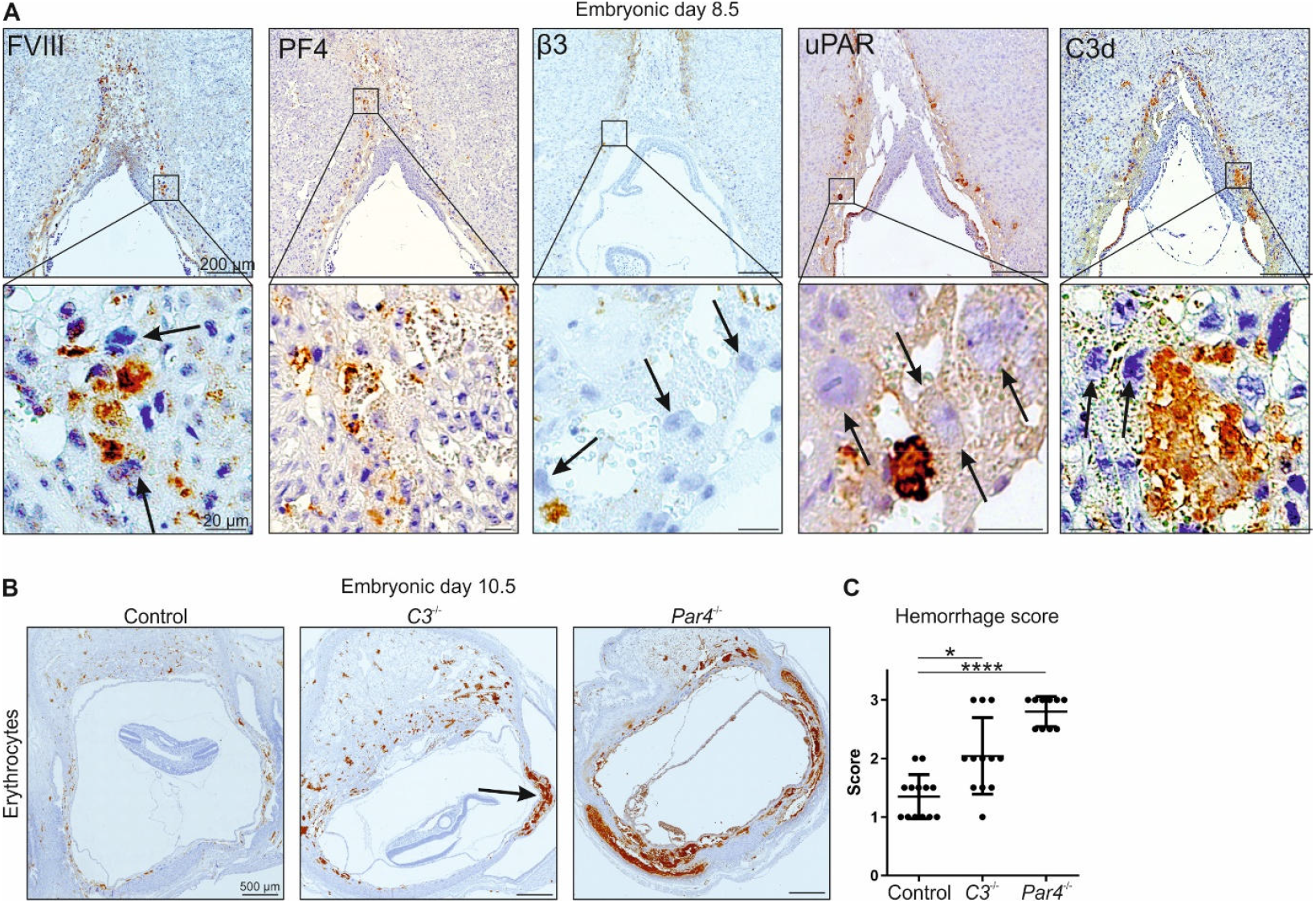
Coagulation and complement regulate maternal blood flow at the developing placenta. (**A**) Representative images of FVIII, PF4, β3, uPAR and C3d immunostaining of placenta sections at E8.5. Arrows point towards polyploid trophoblast giant cells. For the number of individually analysed implants, see Table S1. (**B**) Representative images of distribution of maternal erythrocytes (TER-119) at E10.5 in pregnancies of control, *C3*^-/-^ and *Par4*^-/-^ mice. Arrow points towards antimesometrial haemorrhage. (**C**) Haemorrhage score of (B), ranging from 1 (none to mild), 2 (intermediate) to 3 (severe). Data are presented as mean ± SD. Kruskal-Wallis test ^*^P < 0.05, ^***^P < 0.0001. n=13 for controls from one pregnancy, n=12 for *C3*^-/-^ from 2 pregnancies and n=10 for *Par4*^-/-^ from 2 pregnancies.

We then investigated the influence of platelets and the complement system on the direction of blood flow at the developing placenta. At E10.5, when maternal blood flow through the placenta is generally established, the placenta of control animals showed few maternal erythrocytes at the implantation site near the antimesometrial area, indicating a functional barrier and established blood flow through the placenta (Fig. 1B). In contrast, C3- or PAR4-negative pregnancies showed increased deposition of erythrocytes, indicating failed guidance of the blood flow. Using a score ranging from 1 (mild) to 3 (severe), we found significantly more haemorrhage in C3- and PAR4-negative animals compared to control animals.

### PAR4 mediated platelet activation is crucial for PMT formation

As the absence of both C3 and PAR4 leads to deficits in guidance of maternal blood, we examined the composition of PMTs at the fetal-maternal interface in C3- and PAR4-negative animals at E8.5. In placentas from PAR4-negative mothers, FVIII was no longer enriched in clots, but diffusely associated with TGCs (Fig. 2A). Moreover, only very few platelets (PF4) and hardly any reactivity for C3d were found (Fig. 2A), indicating a complete absence of PMTs. In C3-negative pregnancies, all the aforementioned components were present except for C3 (Fig. 2B). These data imply that PAR4-mediated platelet activation at the fetal-maternal interface is essential for formation of PMTs, which significantly contribute to regulating maternal blood flow at the developing placenta. In the absence of C3, PMT formation was observed but unable to sufficiently guide blood flow to the placenta and to prevent antimesometrial haemorrhage.

**Figure 2:**
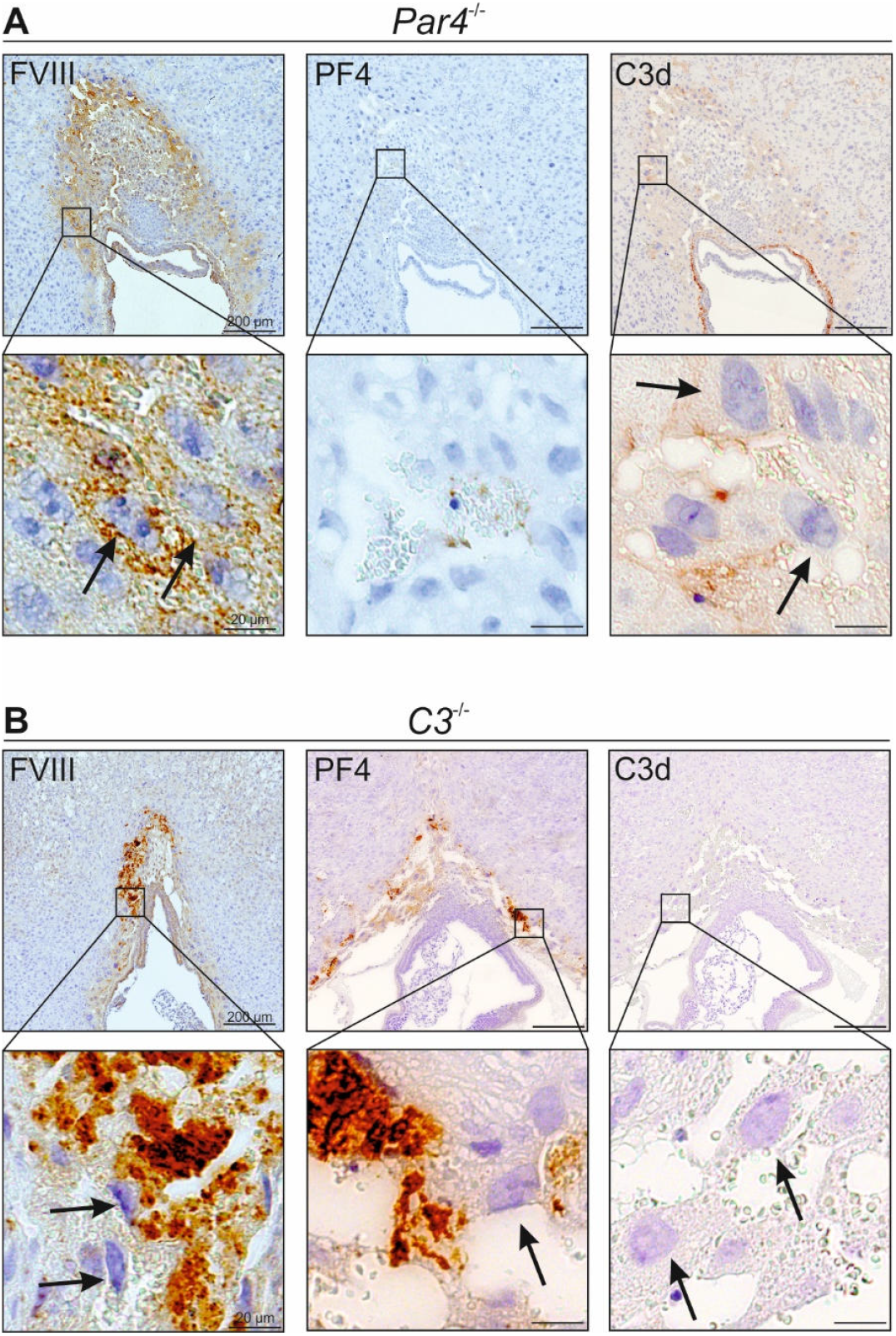
PAR4 mediated platelet activation is crucial for thrombus formation at the placenta. (**A**) Representative images of FVIII, PF4 and C3d immunostaining of *Par4*^-/-^ placenta sections at E8.5. Arrows point towards polyploid trophoblast giant cells. (**B**) Representative images of FVIII, PF4 and C3d immunostaining of *C3*^-/-^ placenta sections at E8.5. Arrows point towards polyploid trophoblast giant cells. For the number of individually analysed implants, see Table S1.

### Loss of placental complement regulation results in excessive thrombosis

Since excessive complement activation is associated with human pregnancy complications and mouse models with impaired complement regulation show severe placental developmental deficits, we asked if platelet activation contributes to complement-mediated placental malformations. As we have previously demonstrated, loss of sialoglycans, due to impaired activation of sialic acid (*Cmas*^−/−^), results in excessive activation of the maternal complement system and pregnancy loss (7) (Fig. 3A). Investigation of coagulation proteins at the fetal-maternal interface of *Cmas*^−/−^ mice revealed that FVIII positive thrombi form around the implantation site with increased numbers and size (Fig. 3B). Likewise, the implantation site of *Cmas*^−/−^ animals revealed a pronounced infiltration with maternal platelets (Fig. 3C-D). To analyse whether this pathological thrombosis depends on activation of the complement system, *Cmas*^−/−^ embryos were analysed on a C3-negative background. C3 deficiency rescued the placental developmental deficit (Fig. S1). Formation of PMTs, deposition of FVIII and maternal platelets were similar to normal controls again (Fig. S1). These data demonstrate that loss of complement regulation can cause excessive thrombosis, which can be corrected by C3 deletion.

**Figure 3:**
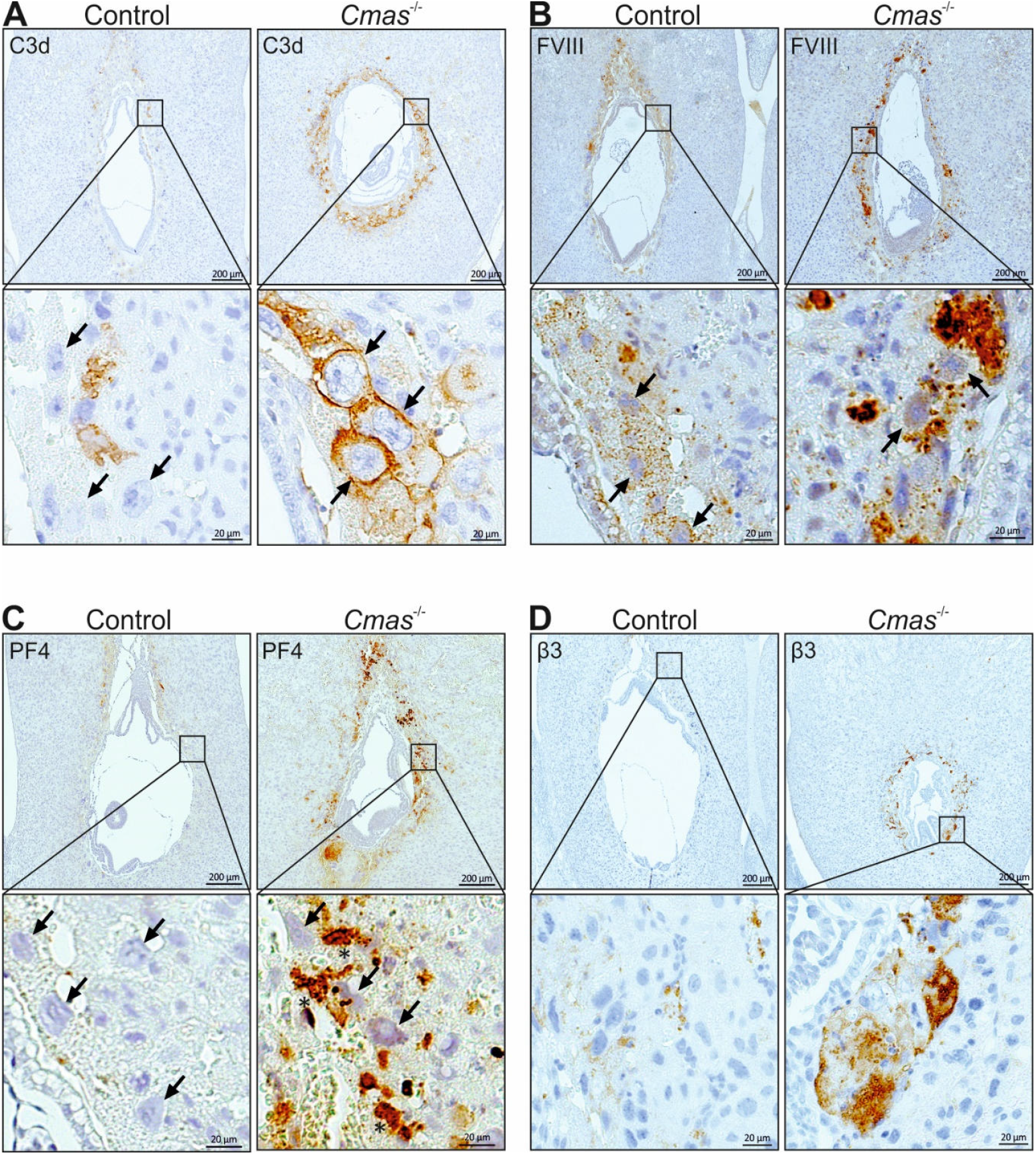
Loss of placental complement regulation results in excessive thrombosis. (**A-D**) Representative images of C3d (**A**), FVIII (**B**), PF4 (**C**) and β3 (**D**) immunostaining of Control and *Cmas*^-/-^ placenta sections at E8.5. Arrows point towards polyploid trophoblast giant cells. For the number of individually analysed implants, see Table S1.

### C5b-9 and anaphylatoxin signaling do not contribute to thrombosis in *Cmas*^-/-^ pregnancies

Formation of the membrane attack complex (MAC or C5b-9) is the terminal step in the complement cascade, and several studies have demonstrated procoagulant properties of C5b-9 (11, 12). However, interrupting the complement cascade already at the step of C5 activation had no impact on excessive thrombosis and pregnancy loss in *Cmas*^-/-^ pregnancies (Fig. S2A-D). Of note, the pronounced mesometrial C3d and PF4 reactivity in controls under C5 blockade indicate that C5b-9 is not involved in PMT formation, further strengthening the point that platelet activation and C3 are the main driver of clot formation at the developing placenta.

We also addressed the potential role of the anaphylatoxins C3a and C5a, as these inflammatory mediators can contribute to a procoagulant environment (13). However, *Cmas*^−/−^ mice showed no improvement in complement activation, thrombosis or placental development on a C3aR/C5aR1-negative background (Fig. S3A-C).

Together, the terminal complement cascade and anaphylatoxin signalling were not involved in excessive thrombosis observed in CMAS-deficient pregnancies. The activation of platelets and C3 were sufficient to cause thrombosis.

### Platelet activation stimulates C3 deposition of Cmas-deficient trophoblasts

As PAR4-mediated platelet activation was required for PMTs and complement activation, the role of PAR4 was also investigated in CMAS-negative pregnancies. Indeed, C3d deposition on the cell surface of CMAS-negative TGCs was absent in PAR4-negative pregnancies (Fig. 4A). However, a diffuse reactivity of C3d could still be observed in the vicinity of these cells. Analysis of the alternative complement pathway stabilizer properdin revealed a very similar picture. Properdin was localized to the cell surface of CMAS-negative TGCs in *Par4*^+/+^, but not in *Par4*^-/-^ pregnancies (Fig. 4B). Instead, diffuse deposition of properdin was observed, similar to C3d, in *Cmas*^-/-^; *Par*4^-/-^ animals. Immunohistochemical analysis for PF4 and β3 showed that *Par*4^-/-^ maternal thrombocytes were still recruited to the fetal-maternal boundary layer in *Par*4^-/-^; *Cmas*^-/-^ pregnancies (Fig. 4C). Although no FVIII-positive thrombi were found, there was generally increased reactivity to FVIII in *Cmas*^-/-^; *Par*4^-/-^ pregnancies (Fig. 4C). Notably, despite the reduced complement activation on the cell surface, there was no improvement in placental development in *Cmas*^-/-^; *Par*4^-/-^ pregnancies as evidenced by the absence of the chorionic plate (CP) (Fig. 4D). It can be concluded that the absence of PAR4-mediated platelet activation prevents the deposition of C3 on Sia-negative cell surfaces; however, *Par4*^-/-^ platelets and coagulation factors are still abundant in the *Cmas*^-/-^ background and placental development remains severely disturbed.

**Figure 4:**
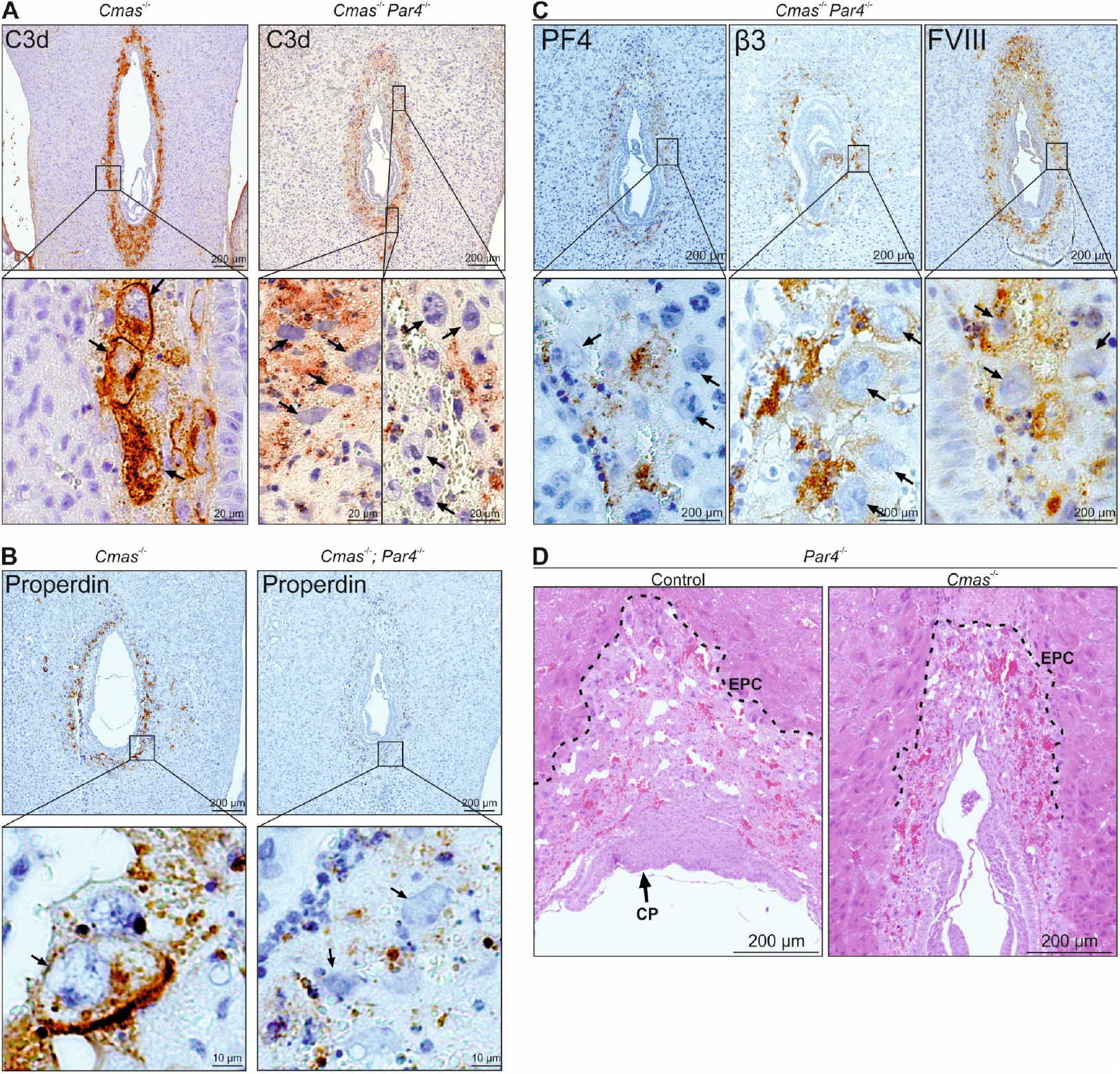
PAR4 mediated platelet activation drives C3 deposition on trophoblasts, but is not solely responsible for platelet recruitment to the fetal-maternal interface. (**A-B**) Representative images of C3d (**A**) and Properdin (**B**) immunostaining of *Cmas*^-/-^ and *Cmas*^-/-^; *Par4*^-/-^ placenta sections at E8.5. Arrows point towards polyploid trophoblast giant cells. (**C**) Representative images of PF4, FVIII and β3 immunostaining of *Cmas*^-/-^; *Par4*^-/-^ placenta sections at E8.5. Arrows point towards polyploid trophoblast giant cells. (**D**) H&E staining of *Cmas*^-/-^ implants at E8.5 in *Par4*^-/-^ mothers. CP, chorionic plate; EPC, ectoplacental cone. For the number of individually analysed implants, see Table S1.

### Depletion of maternal platelets mitigates excessive thrombosis and improves placental development in *Cmas*^-/-^ pregnancies

To investigate this hypothesis further, maternal platelets were depleted in the maternal circulation by anti-GPIbα injections during pregnancy. Platelet depletion abolished deposition of C3d on TGCs and reactivity with properdin (Fig. 5A-B). The diffuse C3d and properdin reactivity still observed in PAR4-negative pregnancies was markedly reduced upon platelet depletion (Fig. 5A-B). Similar to controls (Fig. 1A and 3B) FVIII reactivity in *Cmas*^-/-^ mice was only found in PMTs at the mesometrial side of the fetal-maternal interface (Fig. 5C). Moreover, also development of the placenta was markedly improved in CMAS-negative implants in platelet-depleted mothers (Fig. 5D).

**Figure 5:**
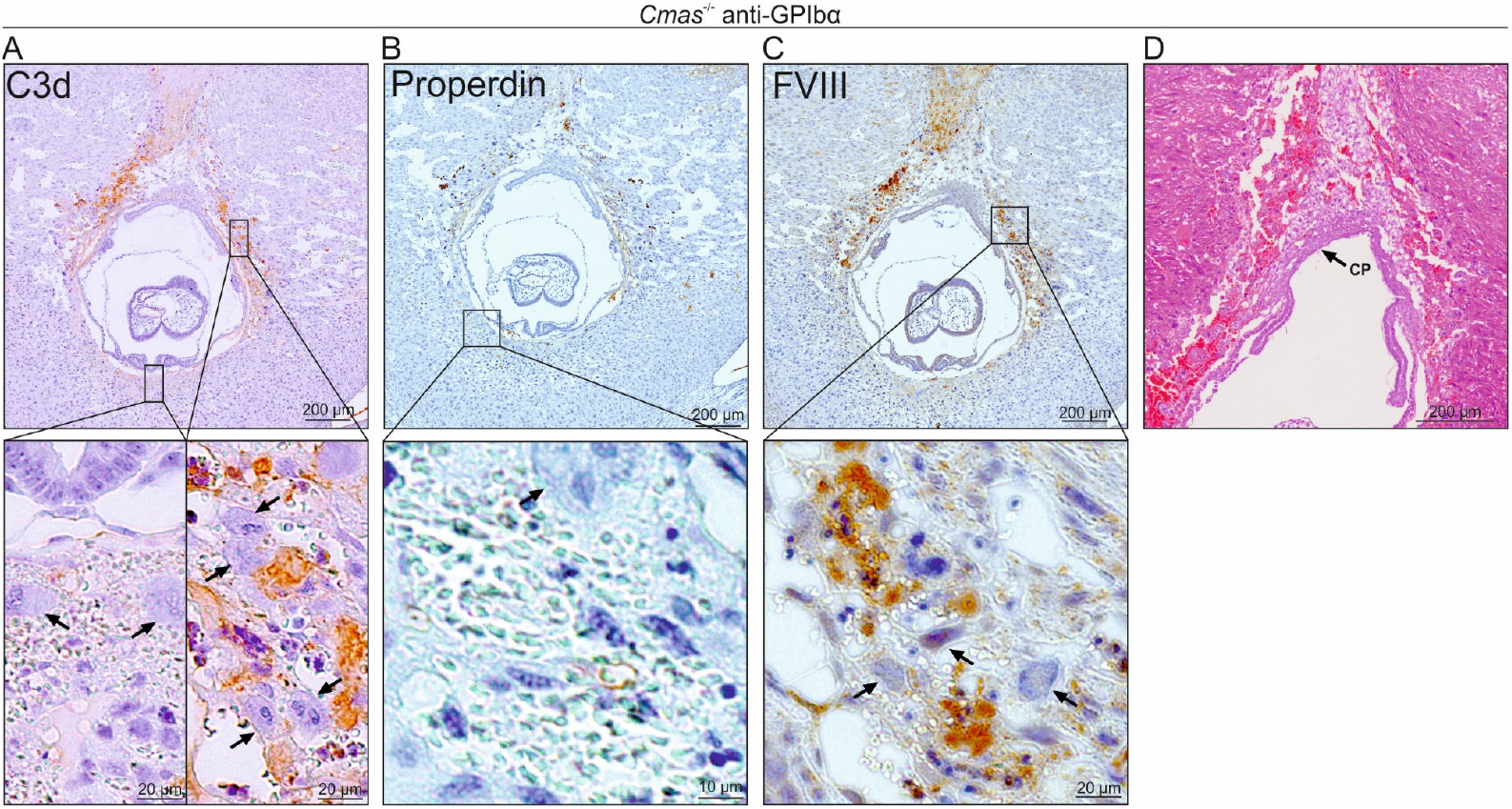
Platelet depletion mitigates excessive thrombosis and developmental deficits of the placenta. (**A-C**) Representative images of C3d (**A**), Properdin (**B**) and FVIII (**C**) immunostaining of *Cmas*^-/-^ implants sections at E8.5 in anti-GPIbα treated mothers. Platelets were depleted by intraperitoneal injection of 200 µg anti-GPIbα (R300 Emfret Analytics) at E2.5 and E5.5 in 3 individual experiments. Arrows point towards polyploid trophoblast giant cells. (**D**) H&E staining of *Cmas*^-/-^ implants at E8.5 in anti-GPIbα treated mothers. CP, chorionic plate. For the number of individually analysed implants, see Table S1.

This data strongly suggests that – despite the importance of PAR4 activation of platelets in the formation of PTMs – platelets provided a surface for ongoing activation of complement and coagulation in the *Cmas*^-/-^ background that was only rescued with platelet depletion.

### PAR4-deficient platelets can still be activated and contribute to thrombosis in *Cmas*^-/-^ pregnancies

The significant improvement in phenotype observed in pregnancies involving platelet depletion, compared to PAR4-negative animals, suggests that platelet activation may still be occurring in the latter. To investigate this, we analyzed the presence of serotonin (5-HT), which is stored in platelet dense granules and released upon activation. In control animals, we found small serotonin-positive particles that were approximately the size of platelets (Fig. 6). These are most likely non-activated platelets in which serotonin is still stored in dense granules. Interestingly, a few TGC nuclei were also positive for serotonin. A very high reactivity for serotonin was observed in *Cmas*^-/-^ embryos along the entire implantation site, suggesting release from activated platelets. A strong presence of serotonin was also still evident at the fetal-maternal interface in *Par4*^-/-^ animals. A completely different picture emerged in platelet-depleted pregnancies. Apart from the serotonin-positive TGC cell nuclei, which were also observed in the control animals, serotonin was virtually absent.

**Figure 6:**
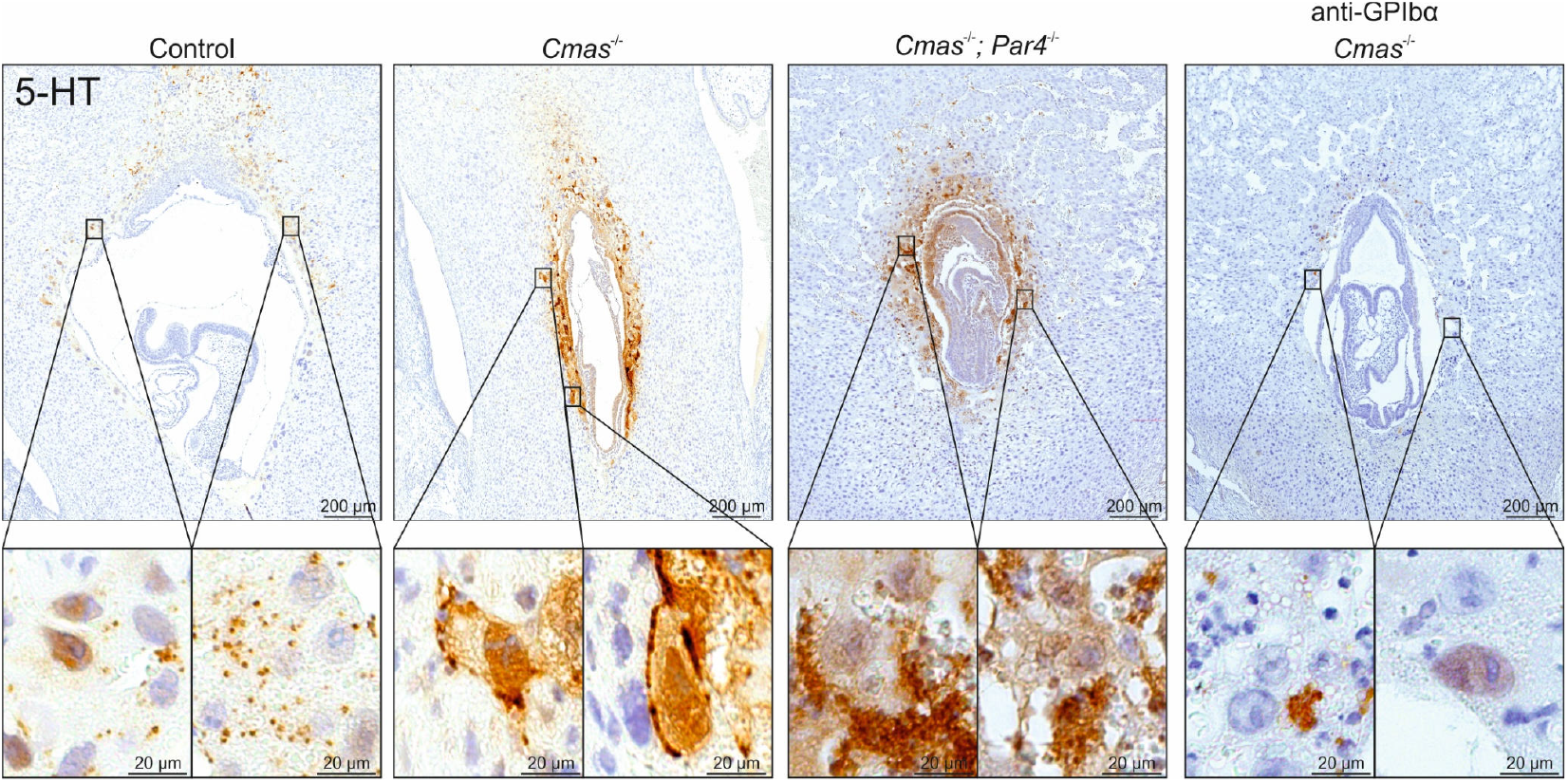
*Par4*^*-/-*^ platelets at the fetal-maternal interface can be activated via thrombin-independent pathways. Representative images of serotonin (5-HT) immunostaining of *Cmas*^-/-^ implants sections at E8.5 in control, *Par4*^-/-^ mothers and anti-GPIbα treated mothers. For the number of individually analysed implants, see Table S1.

## Discussion

In this study, we investigated the regulated formation of placental microthrombi (PMTs), which are essential for directing maternal blood flow into the developing placenta. Our findings demonstrate that deficiencies in complement (*C3*^-/-^) or platelet activation (*Par4*^-/-^) result in excessive placental bleeding. This underscores the necessity of both systems in early pregnancy. Additionally, our findings reveal that PAR4-mediated platelet activation is necessary to localize C3 activation to PMTs, thereby linking coagulation and complement in placental development. Conversely, excessive complement activation in sialoglycan deficient animals results in pathological thrombosis. This phenotype can be reversed by modulating platelet activity.

The role of PAR4 in linking coagulation and complement activation on TGCs is particularly intriguing. Platelet activation through PAR4 occurs upon cleavage of the receptor by thrombin, the central enzyme of the coagulation cascade. Thrombin generation in early pregnancy appears to rely primarily on the extrinsic pathway, since deficiency of tissue factor (TF), factor X, V, or prothrombin results in embryonic lethality around E9-E10 (14-17). In contrast, loss of intrinsic pathway components (including factors VIII, IX, XI) – or von Willebrand factor – does not impair embryonic survival, consistent with their dispensability for placental development. TGCs strongly express TF, and local activation of the TF-FVIIa-FX pathway provides a source of thrombin that activates platelets via PAR4 (4). Once activated, platelets contribute to thrombosis by exposing a procoagulant phospholipid surface, supplying coagulation factors, and releasing granule contents that increase the local calcium concentration. Our findings further demonstrate that PAR4-mediated platelet activation is required for complement C3 activation, and that deficiency of either PAR4 or C3 results in excessive bleeding. Together, these data suggest that TF-driven coagulation on TGCs must be amplified by platelet and complement activation to enable proper PMT formation.

On the other hand, the interplay between coagulation, platelets, and complement requires tight regulation to prevent pathological outcomes. In CMAS deficiency, factor H-mediated inactivation of C3 is impaired, leading to excessive complement activation and thrombosis – a process that can be called *thromboinflammation* – in the developing placenta. This pathological phenotype was partially rescued by either PAR4 deficiency and – even more completely – by maternal platelet depletion, demonstrating that platelets are necessary for C3 deposition and complement-driven thrombosis at the maternal-fetal interface. These findings highlight that, while platelet-driven complement activation is critical for proper PMT formation and placental hemostasis, dysregulation of this crosstalk can rapidly shift the balance toward excessive thromboinflammation. The procoagulant environment of the placenta is also sensitivity to excessive coagulation reactions, which is demonstrated in mice with deficiency in thrombomodulin (TM) or the endothelial protein C receptor (EPCR); these mice die *in utero* around the same time as *Cmas* and *Crry*-negative mice (7, 8).

The mechanisms described in this study are likely relevant to human disease. Platelet-derived C3 has been shown to contribute to inflammation in human influenza virus infection (18), and activated platelets can serve as a surface for complement activation on other cells (19, 20). In systemic lupus erythematosus, platelet activation collaborates with antiphospholipid antibodies to drive complement activation (21). Conversely, complement inhibition rescued a patient with severe coagulation and platelet activation in the context of vaccine-induced thrombotic thrombocytopenia (22). Excessive complement activation has also been linked to pregnancy complications, including preeclampsia (23) and in patients with SLE who experience adverse pregnancy outcomes (24). Genetic polymorphisms in complement regulatory proteins, including factor H, CD46, and C4b-binding protein, which reduce expression or function, are associated with increased risk of severe preeclampsia and recurrent pregnancy loss (25, 26). Proteomic analyses further show that complement and coagulation proteins are highly dysregulated in early-onset severe preeclampsia compared to controls, underscoring the close interplay of these pathways in adverse pregnancy outcomes (27).

Our data suggest that excessive complement activation contributes to pathological thrombosis in the placenta and can be mitigated by C3 deficiency or platelet depletion. Notably, deficiency in PAR4, unlike anti-GPIbα platelet depletion, did not completely prevent C3 activation in CMAS-deficient pregnancies, indicating that platelets activated through other pathways – such as ADP or prostaglandins – can also support complement activation. In the clinical setting, combination therapy with aspirin and low-molecular-weight heparin is the standard of care to prevent pregnancy loss in patients with antiphospholipid antibodies, but it is not always effective (28). Our findings suggest that targeting additional platelet activation pathways, together with inhibition of the alternative complement pathway, may represent a promising strategy for future clinical studies aimed at preventing placental thrombosis and pregnancy loss (29).

While our study provides new insight into the intricate balance of coagulation, platelet, and complement activation during placental development, several limitations should be acknowledged. First, although we demonstrate that platelet activation via PAR4 is required for C3 activation and PMT formation, the precise molecular interactions between TGCs and platelets remain unclear, and the mechanism by which platelets promote complement activation is not fully resolved. Second, we did not directly address the molecular mechanisms by which C3 activation or deposition contributes to placental thromboinflammation. Our data suggest that inhibition of C3a and C5a receptors or downstream complement/MAC activation does not prevent excessive thrombosis. While C3b-dependent amplification of complement and coagulation has been described in other contexts (30), the exact molecular mechanism in the placental environment remains to be elucidated. Third, our findings are based on mouse models, and species-specific differences in platelet and complement biology may limit direct extrapolation to humans. Finally, although we used genetic and depletion approaches to manipulate platelets and complement, these interventions do not fully capture the subtleties of temporal and spatial regulation in normal pregnancy. Future studies using advanced imaging, cell-specific reporters, and human placental models will be required to dissect these interactions in greater detail.

In summary, our study reveals that the tightly coordinated activation of platelets, coagulation, and complement is essential for proper PMT formation and placental development. Disruption of this balance, as seen with PAR4 or CMAS deficiency, leads to pathological bleeding or thrombosis, highlighting the interdependence of these pathways. Our findings provide mechanistic insight into how dysregulated thromboinflammation may contribute to pregnancy complications and suggest that targeted modulation of platelet and complement activity could represent a promising avenue for therapeutic intervention. By illuminating these fundamental processes, our work lays the groundwork for future studies to better understand conditions of adverse pregnancy outcomes and their prevention.

## Supplemental material

**Figure S1:**
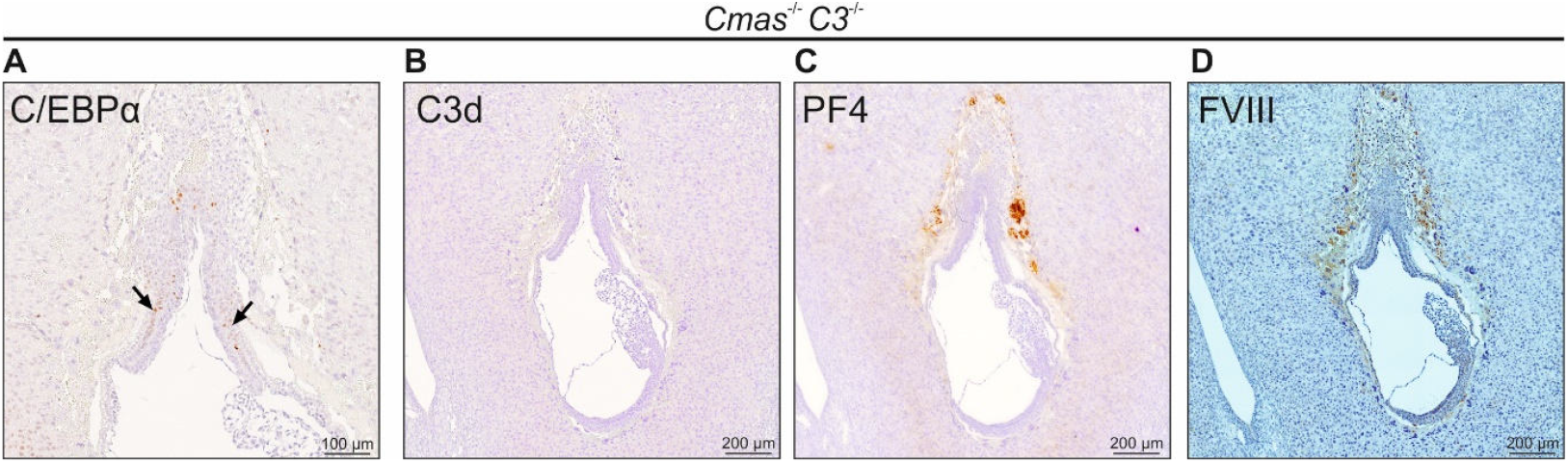
Loss of C3 rescues excessive thrombosis and placental development in *Cmas*^-/-^ mice. (**A-C**) Representative images of C/EBPα (A), C3d (B), PF4 (C) and FVIII (D) immunostaining of *Cmas*^-/-^; *C3*^-/-^ placenta sections at E8.5. Arrows in A point towards the chorionic plate. For the number of individually analysed implants, see Table S1.

**Figure S2:**
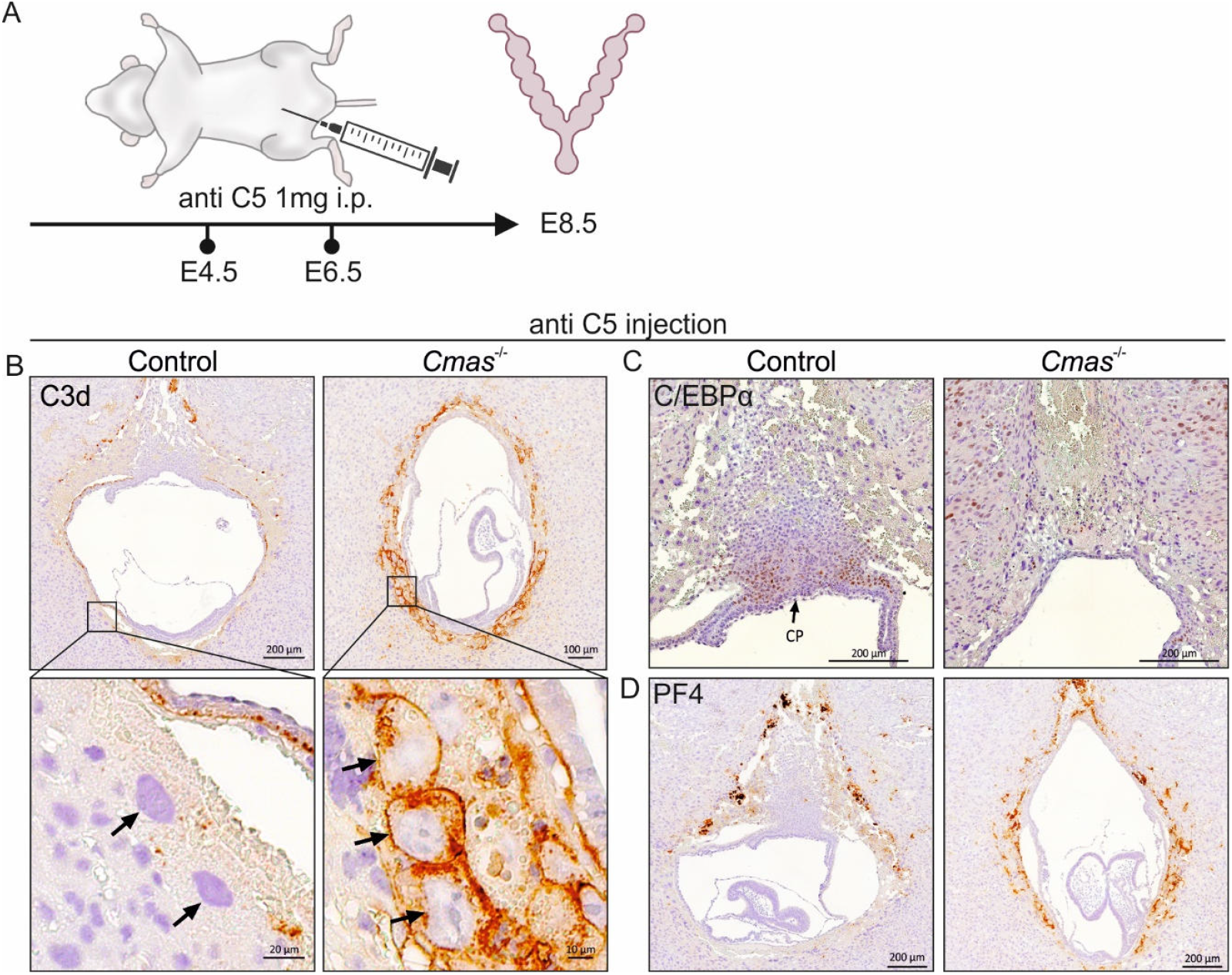
C5 blockade does not reduce excessive thrombosis in *Cmas*^-/-^ implants. (A) Treatment regimen for C5 blockade. C5 activation was blocked by intraperitoneal injection of 1 mg anti-C5 (BB5.1) at E4.5 and E6.5 in 4 individual experiments. (**B-D**) Representative images of C3d (B), C/EBPα (C) and PF4 (D) immunostaining of *Cmas*^-/-^ pregnancies under C5 blockade at E8.5. For the number of individually analysed implants, see Table S1.

**Figure S3:**
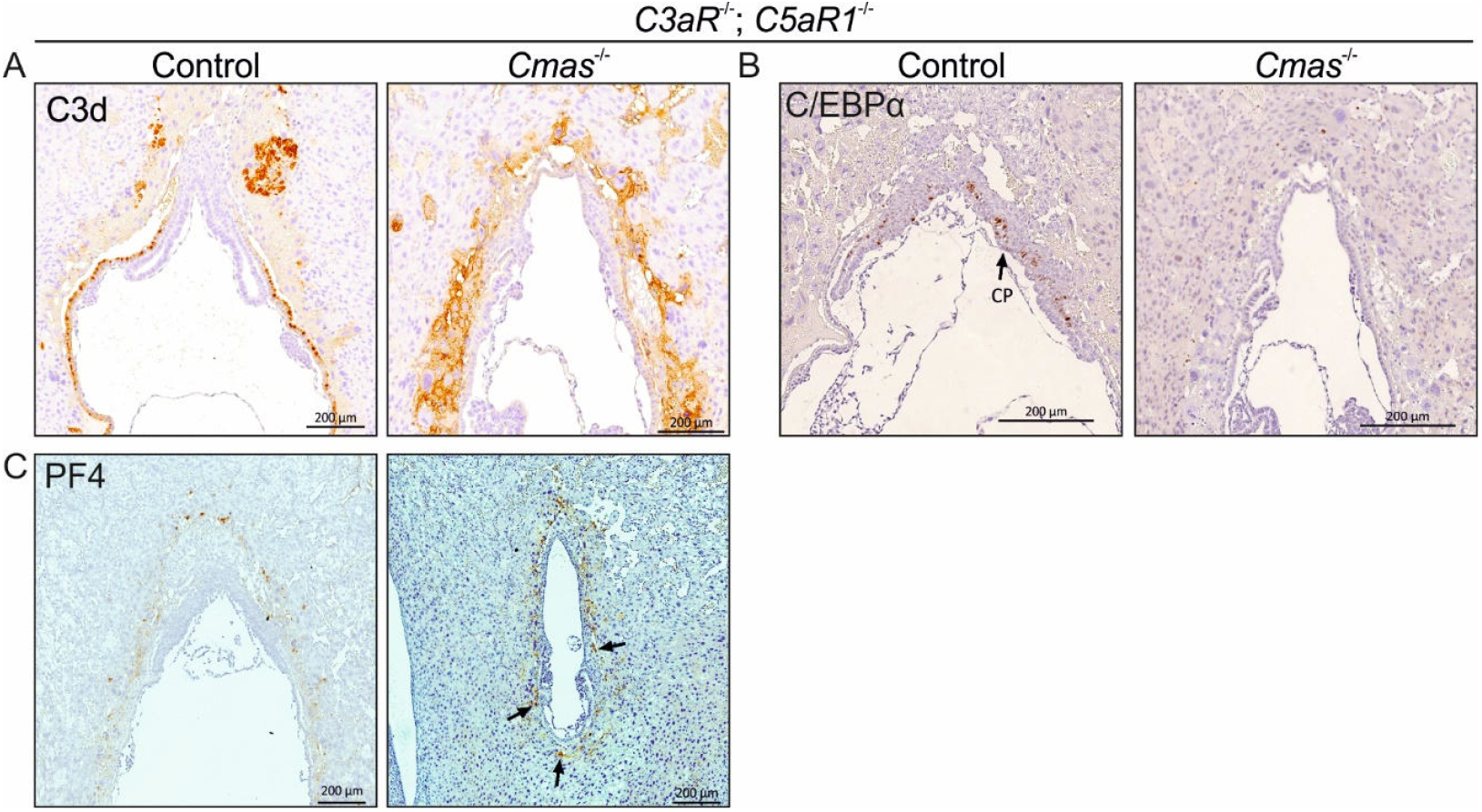
Impaired anaphylatoxin signaling does not rescue the phenotype of *Cmas*^-/-^ mice. (**A-C**) Representative images of C3d (A), C/EBPα (B) and PF4 (C) immunostaining of *Cmas*^-/-^; *C3aR*^-/-^; *C5aR1*^-/-^ pregnancies under C5 blockade at E8.5. For the number of individually analysed implants, see Table S1.

**Table S1:**
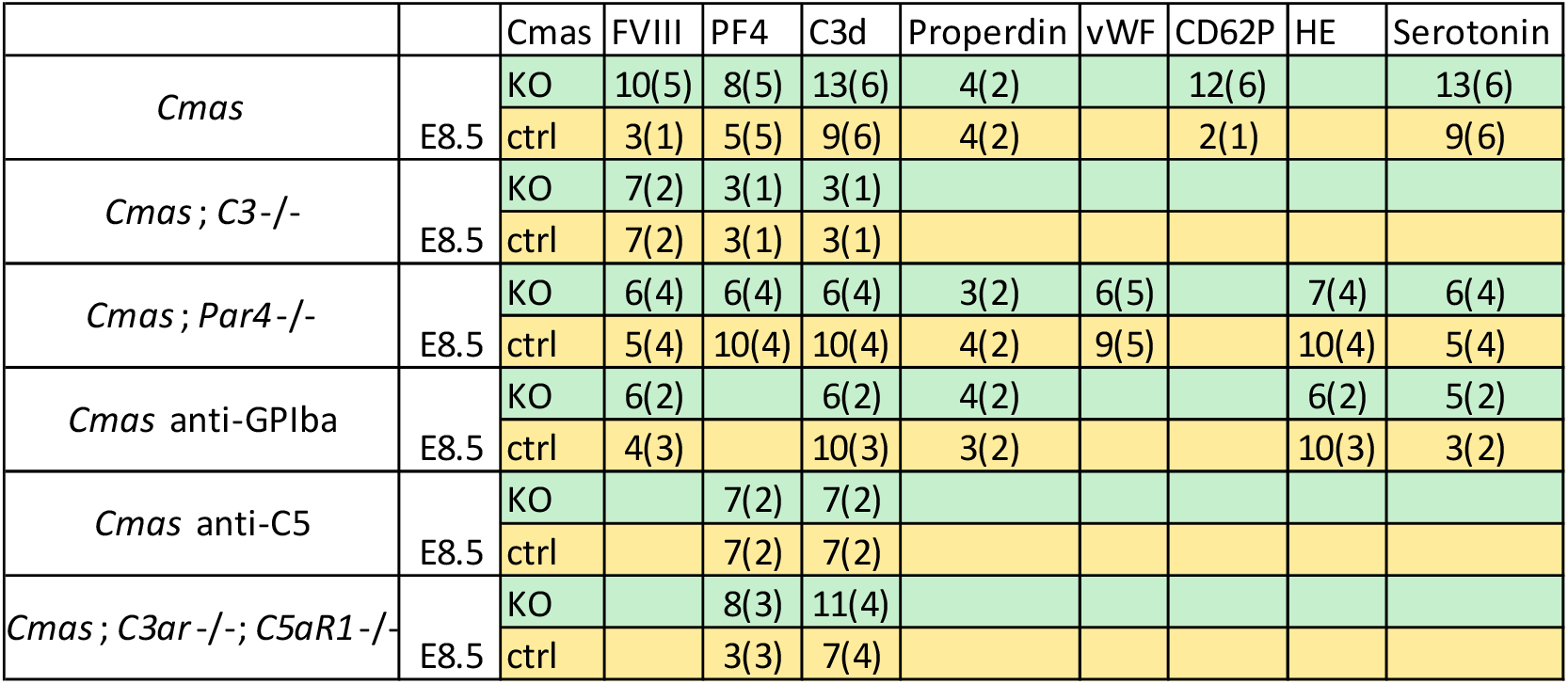
Total numbers of analysed implants and uteri (brackets) in IHC and HE staining.

**Table S2:**
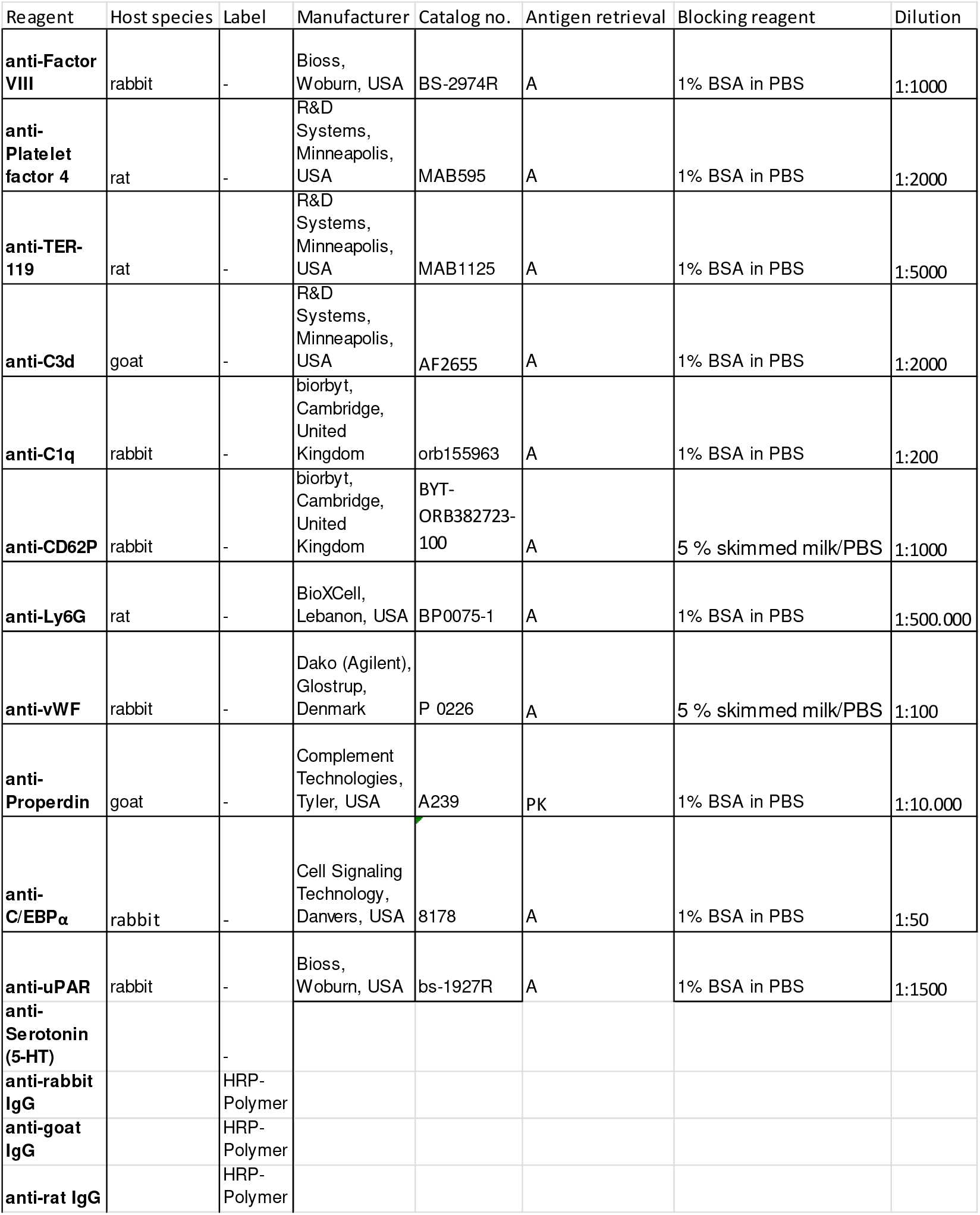
Primary and secondary antibodies used in IHC experiments.

## Methods

### Mice

*Cmas*^*-/-*^ mice (7, 31) were backcrossed for six generations on a NMRI background.*Cmas*^*-/-*^ embryos were compared to *Cmas*^*+/+*^ or *Cmas*^*+/-*^ littermates, referred to as controls.

*Cmas*^*+/-*^; *C3*^*-/-*^ mice were generated by crossing *Cmas*^*+/-*^ animals with the C3-depleted strain *B6*;129S4-C3^tm1Crr^/J. The *B6*;129S4-C3^tm1Crr^/J mice. Upon 4 backcrosses with NMRI mice, heterozygous *Cmas*-knockout mice on a homozygous *C3*-knockout background (*Cmas*^*+/-*^; *C3*^*-/-*^) were obtained (*STOCK*-*C3*^*tm1Crr*^*-Cmas*^*tm3*^). The following primers were used: ms C3geno_for (5′-ATCTTGAGTGCACCAAGCC-3′), ms C3wt_rev (5′-GGTTGCAGCAGTCTATGAAGG-3’) and ms C3mt_rev (5’-GCCAGAGGCCACTTGTGTAG-3’); the PCR program included 3 steps (98°C for 15 seconds, 64,7°C for 30 seconds, 72°C for 30 seconds) and 32 cycles.

*Cmas*^*+/-*^; *C3aR*^*-/-*^; *C5aR1*^*-/-*^ mice were generated by crossing *Cmas*^*+/-*^ animals with the C3aR- and C5aR1-depleted strain *B6*-*C3ar1*^*tm1Raw*^*-C5ar1*^*tm1Cge*^. The *B6*-*C3ar1*^*tm1Raw*^*-C5ar1*^*tm1Cge*^ mice were kindly provided by the Klos Lab (Hannover, Germany). Upon 4 backcrosses with NMRI mice, heterozygous *Cmas*-knockout mice on a homozygous *C3aR; C5aR1*-knockout background (*Cmas*^*+/-*^; *C3aR*^*-/-*^; *C5aR1*^*-/-*^) were obtained (*STOCK C3ar*^*tm1Raw*^*-C5ar*^*tm1Cge*^*Cmas*^*tm3mhhTg15*^). The following primers were used: EW33 (5′-TACAATATAGTCAGTTGGAAGTCAGCC-3′), EW34 (5′-TGGGCTCTATGGCTTCTGAGGCGGAAAG-3’) and EW35 (5’-GAGAATCAGGTGAGCCAAGGAGAA-3’); the PCR program included 3 steps (98°C for 15 seconds, 64,3°C for 30 seconds, 72°C for 30 seconds) and 35 cycles. *C5aR1* PCR: The following primers were used: RL8 (5′-GGTCTCTCCCCAGCATCATA′), RL9 (5′-GGCAACGTAGCCAAGAAAAA-3’) and RL10 (5’-GCCAGAGGCCACTTGTGTAG-3’); the PCR program included 3 steps (98°C for 15 seconds, 60,1°C for 30 seconds, 72°C for 30 seconds) and 35 cycles.

*Cmas*^*+/-*^; *F2rl3*^*-/-*^ mice were generated by crossing *Cmas*^*+/-*^ with the PAR4-depleted strain *B6*.*129S4(FVB)-F2rl3*^*tm1*.*1Cgh*^*/Mmnc*. The *B6*.*129S4(FVB)-F2rl3*^*tm1*.*1Cgh*^*/Mmnc* strain (32) was obtained from the Mutant Mouse Resource and Research Center (MMRRC) at University of North Carolina at Chapel Hill, an NIH-funded strain repository, and was donated to the MMRRC by Shaun Coughlin, M.D., Ph.D., University of California, San Francisco. Heterozygous *Cmas*-knockout mice on a homozygous *F2rl3*-knockout background (*Cmas*^*+/-*^; *F2rl3*^*-/-*^) were obtained (*STOCK F2rl3*^*tm*.*1*.*1Cgh*^*Cmas*^*tm2MhhTg15D/Bwei*^). The following primers were used: Par4 for (5′-CAGATGTTTCCTGGGCTGGGTG-3′), lacZ WT rev (5′-ATTGTGGGTGCCTCAGTGTCCC-3’) and Par4 KO rev (5’-CAGGGTTTTCCCAGTCACGACG-3’); the PCR program included 3 steps (98°C for 15 seconds, 69°C for 30 seconds, 72°C for 15 seconds) and 30 cycles.

Animals were hosted in the animal facility of the Hannover Medical School under specific pathogen–free conditions.

### Animal experiments

Anti-C5 (BB5.1) antibody was produced as previously described (33) and applied at E4.5 and E6.5 via i.p. injection of 1000 µg antibody in 100 µl PBS. At E8.5, females were sacrificed. The uterus was isolated for further processing.

Anti-GPIbα antibody (R300, Emfret Analytics) was applied at E2.5 and E5.5 via i.p. injection of 200 µg antibody in 100 µl PBS. At E8.5, females were sacrificed. The uterus was isolated for further processing.

All animal experiments were approved by the Niedersaechsisches Landesamt fuer Verbraucherschutz und Lebensmittelsicherheit (LAVES), Postfach 3949, Oldenburg 26029, Germany (AZ33.9-42502-04-19/3150; AZ33.12-42502-04-20/3510; AZ2019/244.) and by the Institut fuer Versuchstierkunde und Zentrales Tierlaboratorium, Hannover Medical School, CarlNeuberg-Str. 1, Hannover 30625, Germany.

### Histology

Female mice from heterozygous matings were examined daily in the early morning for the presence of a vaginal plug. The time at which the plug was discovered was considered to be day 0.5 after conception. On days 8.5 and 10.5 of gestation, the pregnant mice were sacrificed and their uteri dissected and fixed in 4% paraformaldehyde in PBS at 4°C for 48 hours. Following fixation, the uteri were dehydrated in a graded ethanol series and embedded in paraffin wax. For histological analysis, the uteri were sectioned into 3 µm slices using a microtome, rehydrated and stained with haematoxylin and eosin (H&E). The slices were analysed using a Zeiss Observer Z1 microscope equipped with a Zeiss AxioCam MRc camera.

### Immunohistochemistry

The deparaffinised tissue was rehydrated and subjected to antigen retrieval using either a target retrieval solution (pH 6, Dako Agilent, Santa Clara, CA, USA) or a proteinase K solution (50 mM Tris (pH 8.5), 1 mM EDTA and 10 µg/ml proteinase K) for 10 minutes at 37 °C. Endogenous peroxidase activity was blocked by incubating the sections in a 3% hydrogen peroxide solution in PBS at room temperature for 30 minutes. This was followed by a blocking step in 1% BSA in PBS for 30 minutes. Primary antibody incubation was performed in the blocking solution overnight at 4 °C. This was followed by a 60-minute incubation with the secondary antibody at room temperature. The primary and secondary antibodies used, as well as the dilutions applied, are listed in table S2. All HRP-conjugated reagents were detected using a DAB (Dako) reaction and were subsequently counterstained with haematoxylin and analysed using the aforementioned microscope setup.

## Author contribution

Designing research studies: A.F., A.T., M.A.

Conducting experiments: A.F., L.S., E.A., M.A.

Acquiring data: A.F., L.S., U.P.B., K.F.S.,

Analyzing data: A.F., L.S., M.A.

Providing reagents: A.K., W.M.Z., B.P.M., A.T.

Writing the manuscript: A.F., A.T., M.A.

Reviewing and editing the manuscript: W.M.Z., B.P.M., A.K.,

## Acknowledgements

We would like to thank Professor Rita Gerardy-Schahn and Dr. Anja Münster-Kühnel for valuable support and productive discussions.

